# Validation of a Single *Nepl15* Transcript in Oregon-R *Drosophila melanogaster* with Minor Coding Sequence Variation Relative to FlyBase

**DOI:** 10.64898/2026.02.20.707127

**Authors:** Chase Drucker, Surya Banerjee

**Affiliations:** Department of Biological Sciences, Texas Tech University, Lubbock, Texas, United States

## Abstract

The *Neprilysin-like 15 (Nepl15)* transcript in *Drosophila melanogaster* displays sex and organ specific phenotypes and only has one known transcript in the fly database (FlyBase.org). Given that the *Nepl15* gene is differentially expressed in a tissue-specific and sex-specific manner, we sought to identify if there were additional *Nepl15* transcripts available in Oregon-R strain flies which had not been reported in the fly database by performing sequencing-based approaches. We have identified presence of different codons in the transcript different than what had been reported in FlyBase. Further experimentation is needed to determine the full effect of these changes on fly physiology.

## Description

Neprilysins (neutral endopeptidases; Neps) belong to the M13 family of Zinc (II) dependent metallopeptidases and was first discovered in mammals (Turner et al., 2001). Previous homology studies utilizing this mammalian protein sequence have identified the presence of Nep, or ECE-like orthologs, in invertebrate organisms including *Drosophila melanogaster* (Bland et al., 2008). Typical mammalian Neps, including Neps found in humans (HsNep), have a short cytoplasmic N-terminal domain, a shorter hydrophobic transmembrane domain, and a long extracellular C-terminal domain that houses highly conserved catalytic motifs. Two of these motifs, HEXXH and EXXAD, are responsible for Zn (II) binding and the subsequent catalytic activity. Two additional motifs, NAY/FY and CXXW, contribute to substrate binding and proteostasis, respectively. This structural arrangement enables Zn (II)-dependent catalytic activity, which in turn guides proteolytic cleavage, typically occurring at the N-terminal side of a hydrophobic residue with a strong preference for Phe or Leu toward the C-terminus (Nalivaeva & Turner, 2013; Sitnik et al., 2014; Turner et al., 2000).

Typical mammalian Neps are normally expressed in a variety of cell types including brush border cells of kidneys, brain, neutrophils, lungs, synapses, along with adipocytes and smooth muscle cells present throughout gut tissue to name a few. Once cleaved, they typically control signals that impact nervous, cardiovascular, inflammatory and immune systems (Schling & Schäfer, 2002; Turner et al., 2001). Interestingly, recent investigation into Nep suggests that it may have a role in blood pressure homeostasis and lipid metabolism. In humans, elevated plasma Nep levels are positively correlated with features of metabolic syndrome (MetS), including BMI, lipid metabolism, and insulin resistance, all of which are associated with elevated cardiometabolic risk. Similarly, high-fat diet-fed mice models exhibit increased Nep expression and activity with obesity, mirroring patterns observed in humans with metabolic syndrome. However, in obesity-induced mice, circulating glucose levels, glucose uptake, and average body weight remain elevated despite Nep inhibition (Standeven et al., 2011). Yet, the potential therapeutic benefits of neprilysin inhibition (specifically, angiotensin receptor– neprilysin inhibition) in patients with heart failure and reduced ejection fraction have been reinforced by clinical trials (Jessup, 2014; Solomon et al., 2019). Collectively, these findings suggest that Nep-associated regulation of glucose homeostasis may arise from conserved molecular pathways rather than organism-specific metabolic effects (Banerjee, 2019).

*Drosophila melanogaster* contain 28 identified neprilysin and neprilysin-like genes. Among them, *Neprilysin-like 15 (Nepl15)*, a predicted secreted, catalytically inactive Nep, transcriptionally rich in fly fat body, regulates glycogen and lipid storage in a sex-dependent manner (Banerjee et al., 2021; Meyer et al., 2021; Turner et al., 2001). Previous work with *Nepl15*^*ko*^ (knock-out) flies revealed that glycogen and triacylglyceride (TAG) storage was significantly reduced in adult male mutants while significantly more carbohydrates and a slightly reduced lipid concentration was observed in adult female mutants (Banerjee et al., 2021). Furthermore, Phosphorous-32 (^32^P) radioisotope labeling of normal food showed no significant difference in the amount of food intake by wild-type (*w*^*1118*^) and *Nepl15*^*ko*^ (*w*^*1118*^; *Nepl15*^*ko*^) adult flies (Banerjee, 2019).

Interestingly, the *Nepl15* transcript is differentially expressed in both larval organs and is expressed 4.5 times more in adult male flies compared to adult female flies. Yet reports from the fly database (FlyBase.org) indicate there is only one *Nepl15* transcript sequence and no intronic regions of this gene in the genome of *y*^*1*^; *cn*^*1*^; *bw*^*1*^; *sp*^*1*^, or ISO1 MT (dm6) strain *D. melanogaster*.

Prior to experimentation, we further investigated and listed all the transcript variants of *D. melanogaster* Neps and Nep-like genes as denoted in FlyBase (release FB2024_02). We found that out of 7 Neps, 6 Neps have multiple transcript variants, of which 3 Neps have more than one unique polypeptide sequence. We also found that out of 21 Nep-like genes, 17 have only one transcript including *Nepl15*, with only 1 of these resulting in more than one unique polypeptide sequence (Table 1).

**Table 1.**
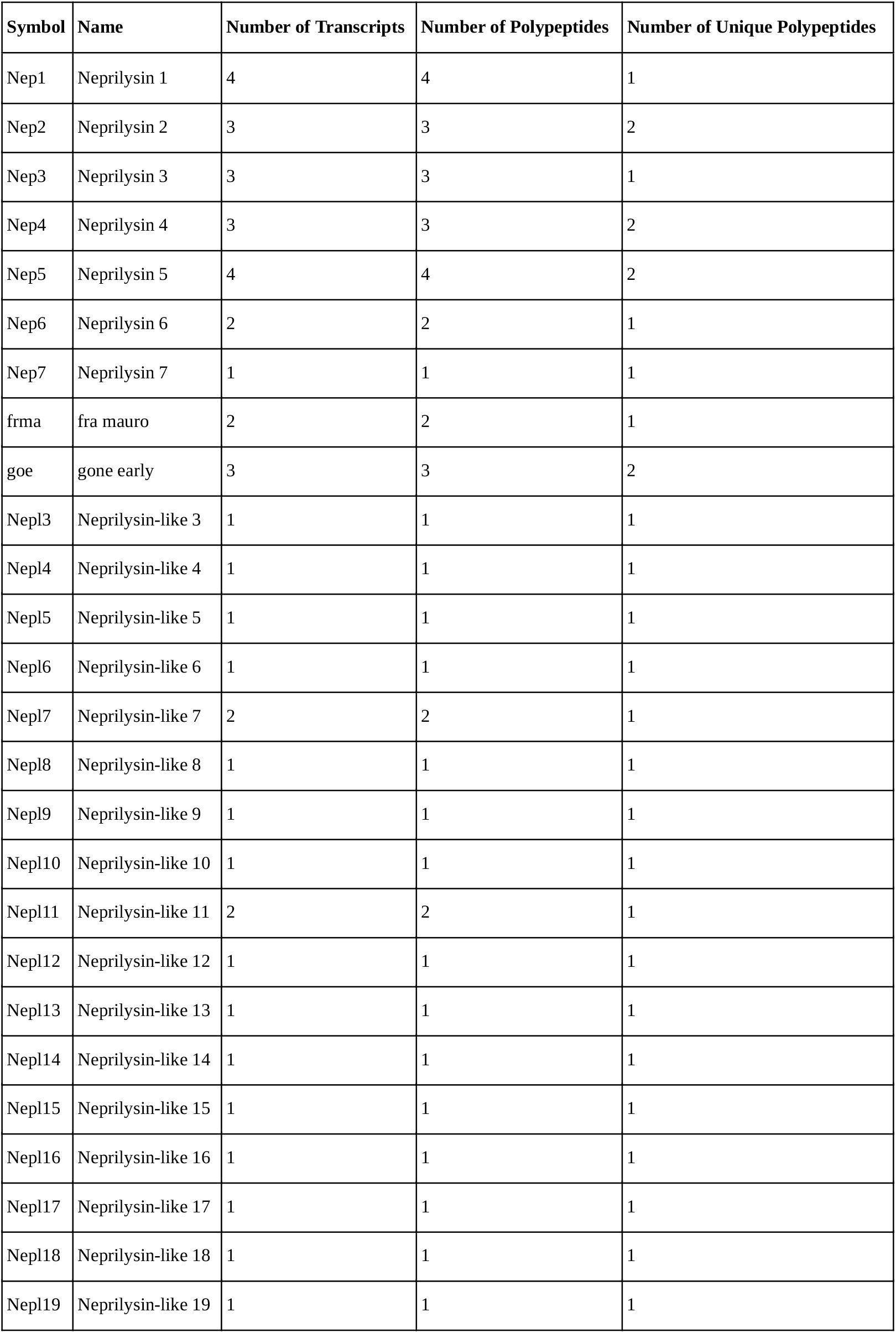
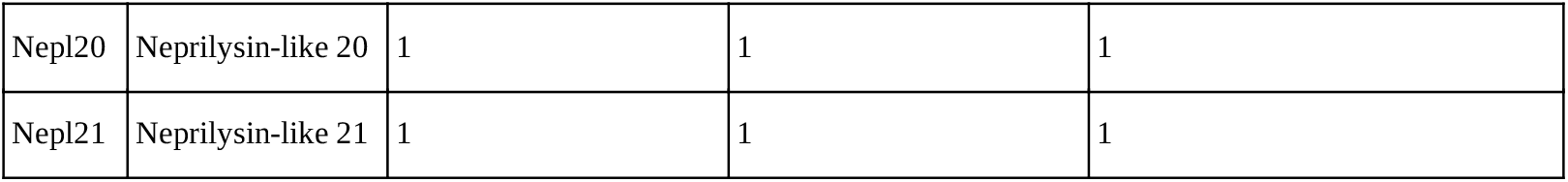
List of Neprilysins and the corresponding number of transcripts and unique polypeptides they produce.

Further analysis of this information led to the formation a couple of major questions which may explain *Nepl15*’s differential gene expression in a tissue and sex-specific manner: (1) Are there any other transcript variants of *Nepl15* in adult male and/or female flies available which have not been recorded? (2) Are there any differences in the 5’ and 3’ UTRs which may regulate differential translation? (3) Are there any differences in the coding sequence?

To answer these questions, we used total RNA from whole-body adult male and female Oregon-R strain flies (FBsn0000276) and performed sequential RT-PCR, PCR, and sequencing-based approaches to identify the presence of a few single nucleotide polymorphisms (SNPs) in the transcript, different than what had been reported in FlyBase. While some of the identified SNPs resulted from missense mutations others resulted from silent mutations. The identified missense mutations were investigated using both TMHMM – 2.0, and SignalP – 6.0, and indicated no differences in protein localization or cleavage site, respectively. Additional analysis utilizing AlphaFold 3 (https://deepmind.google/science/alphafold/; retrieved 6/29/2025) (Abramson et al., 2024) to generate a 3D model of this altered protein and ChimeraX (https://www.rbvi.ucsf.edu/chimerax/; retrieved 6/29/2025) (Meng et al., 2023) to align the predicted 3D structures also indicated minimal alterations in the protein structure (Figure 1. B-C_3_). Further data analysis using Ensembl VEP (https://useast.ensembl.org/Tools/VEP; retrieved 1/16/2026) (McLaren et al., 2016) to further predict the effects of these mutations is summarized in table 2.

**Table 2.**
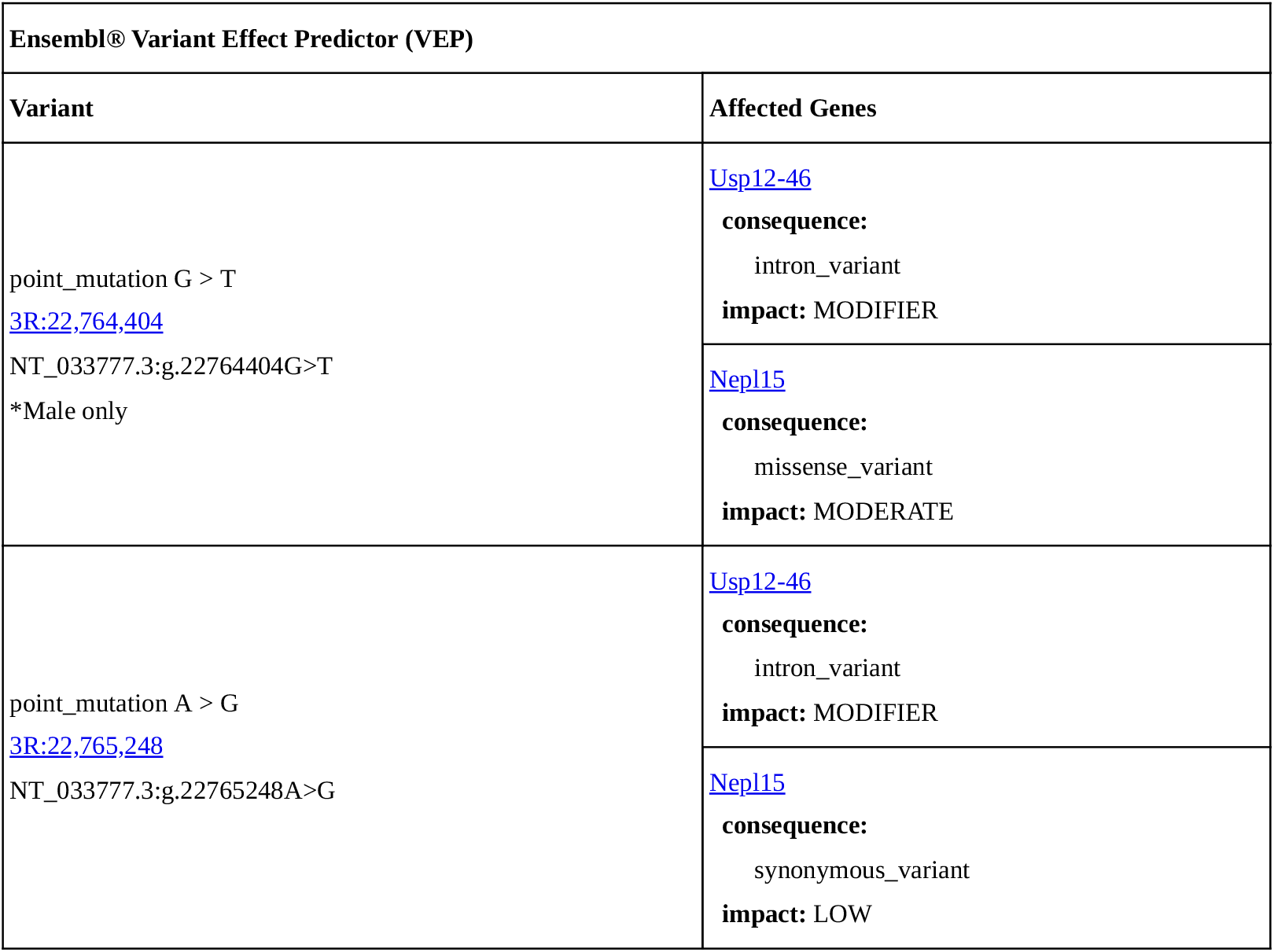
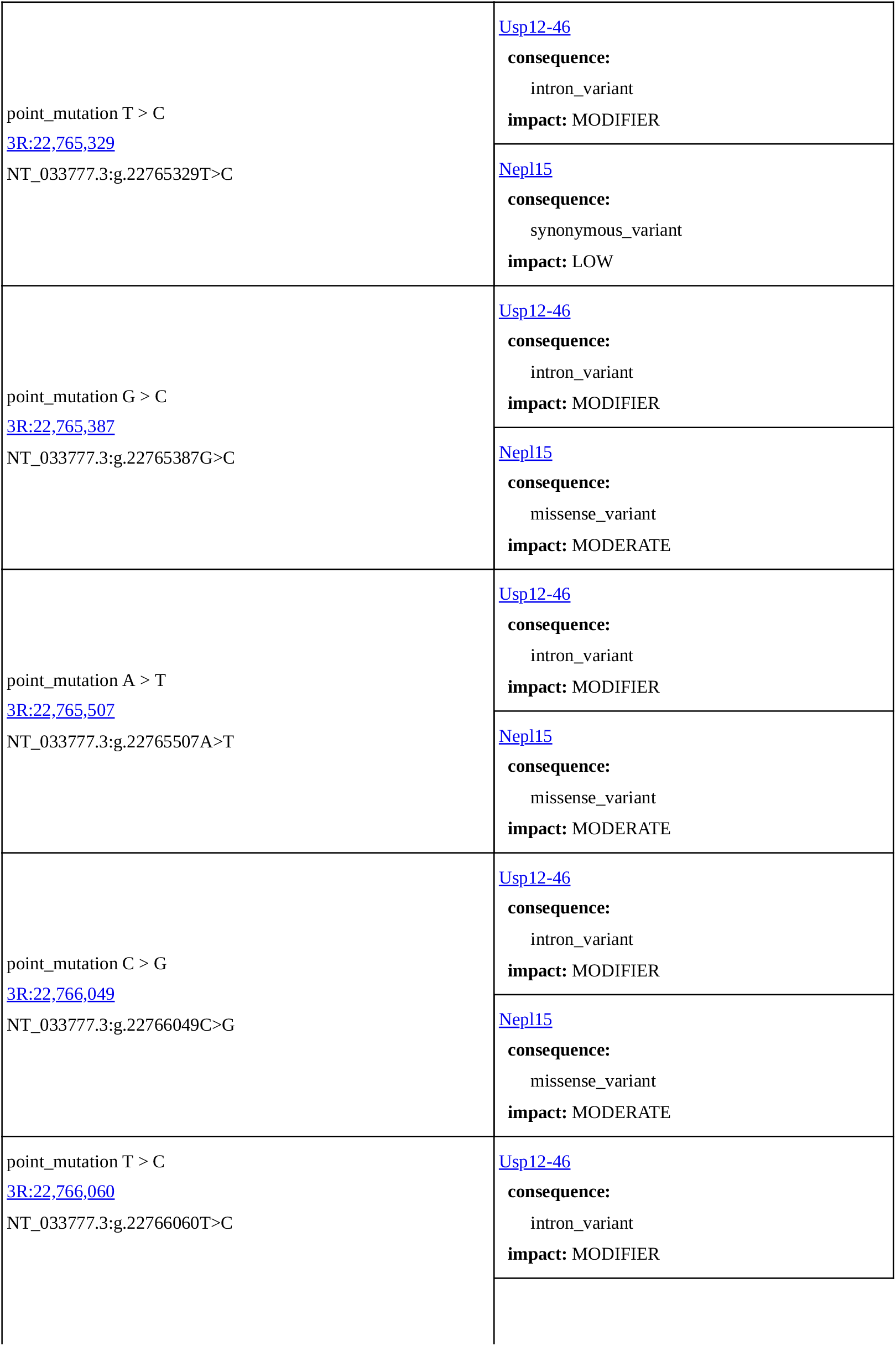
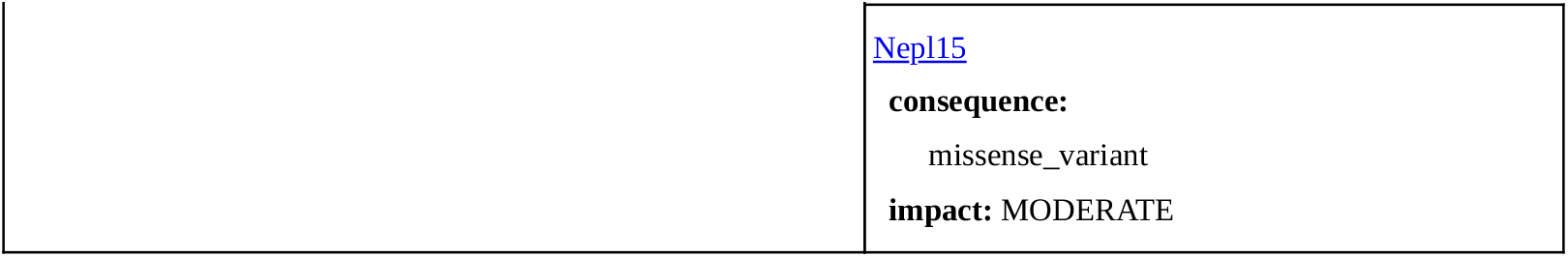
Sequencing alterations and predicted effects observed, according to Ensembl® Variant Effect Predictor (VEP), of both male and female Oregon-R Drosophila melanogaster compared to the Flybase.org sequence.

**Figure 1.**
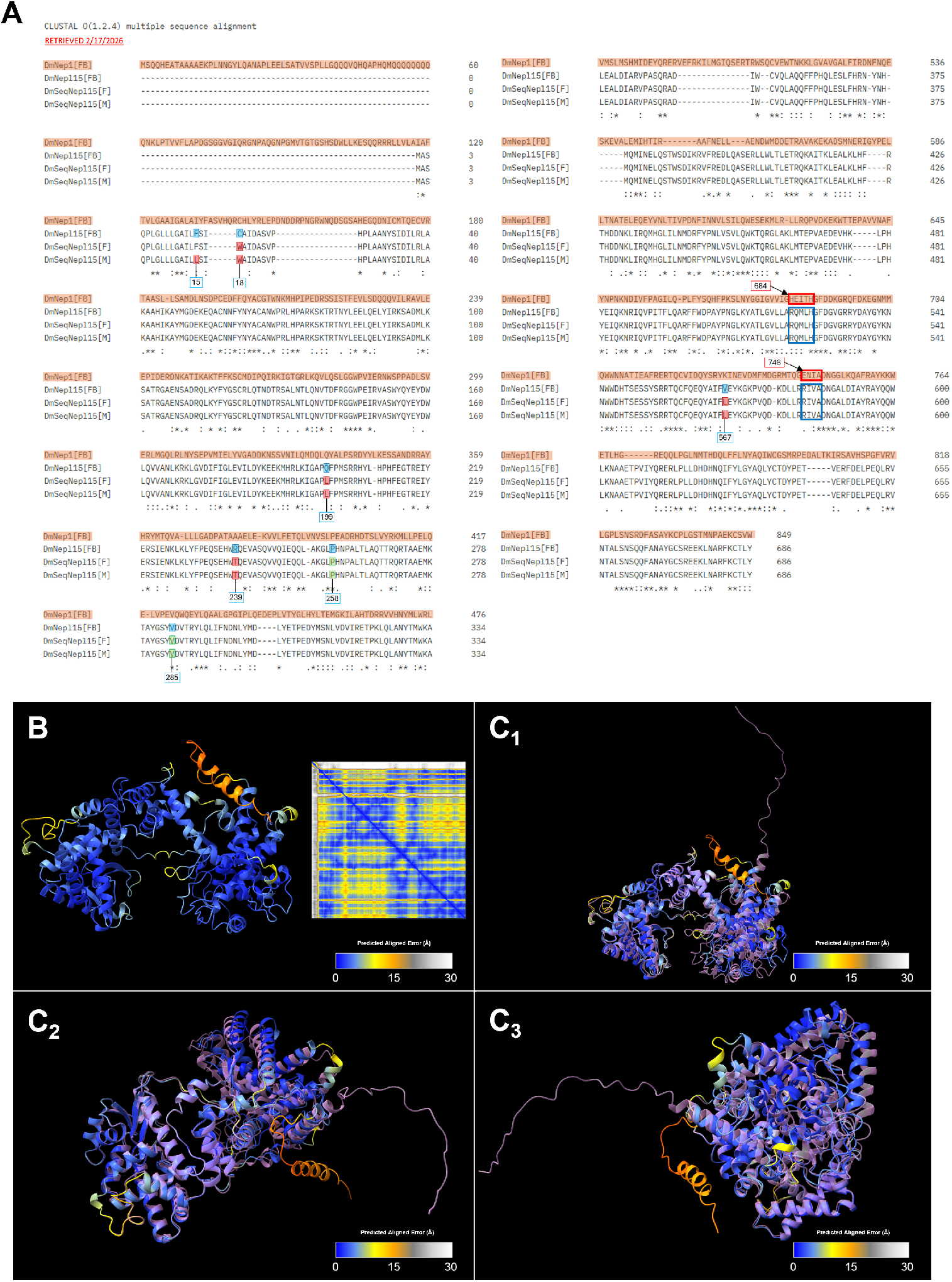
Clustal Omega results for the multiple sequence alignment of sequenced polypeptides and 3D protein prediction and alignment of *Nepl15* peptides: **(A) Clustal Omega results for the multiple sequence alignment of the Nep1 and Nepl15 polypeptide sequences from FlyBase.org [FB] and sequenced polypeptide data from Nepl15 from male [M] and female [F] samples**. Motifs indicated at amino acid positions 684 and 746 indicate that the lack of catalytically conserved residues, HEXXH and EXXA, is conserved in all DmNepl15 sequences. Colored boxes indicate sequenced differences in the polypeptide sequence: (blue) indicating the original residue, (green) designating the locations of silent mutations, and (red) representing missense mutations. The numbers underneath each colored residue correspond to the amino acid position on the Nepl15 sequence. **(B) Predicted 3D AlphaFold model of male Nepl15 sequenced polypeptide**. Right side shows predicted aligned errors (PAE) plot, mapping each expected position error (Å) for pairs of residues, generated by AlphaFold 3. 3D model residues and PAE plot are colored and indicated in the provided key in which lower values (blue) indicate higher levels of confidence. **(C**_**1-3**_**) Predicted 3D model of male Nepl15 sequenced polypeptide aligned with predicted 3D protein structure from FlyBase/UniProt**. All images display different angles of the Nepl15 predicted protein from (B) aligned to the Nepl15 reference polypeptide structure from FlyBase and UniProt databases (light pink).

Collectively, these results seem to indicate that a slightly different *Nepl15* protein is present in our lab grown wild-type Oregon R flies, however the effects of these mutations have yet to be fully experimentally determined. Thus, these results stand to confirm the presence of one *Nepl15* transcript version present in *D. melanogaster* and indicate that our exact sequence seems to differ slightly from the current database, which may be due to a difference in the sequenced strain.

## Methods

### Drosophila melanogaster husbandry

All *Drosophila melanogaster* with the wild-type Oregon-R (OR) strain were maintained using a molasses formula (Flystuff 66-116 Nutri-Fly®) at 25 °C. Population was controlled via the removal of the parental generations of each subsequent generation formed. Age matched 5-to-7-day-old, third generation, flies were used for subsequent PCR and RNA extraction.

### PCR

RNA was extracted from *D. melanogaster* between the ages of 5-7 days old using TRIzol™ reagent (Sigma-Aldrich-T9424) on ice. cDNA synthesis was then performed for samples using 5x iScript Reaction Mix (Bio-Rad-1708889) using the Bio-Rad iScript™ cDNA Synthesis Protocol. Primers were then synthesized using the CDS (coding sequence) and Transcript (T.SCRPT) sequences found on FlyBase (FB) (Table 3). Primers were checked for their efficiency and fidelity before use.

**Table 3.**
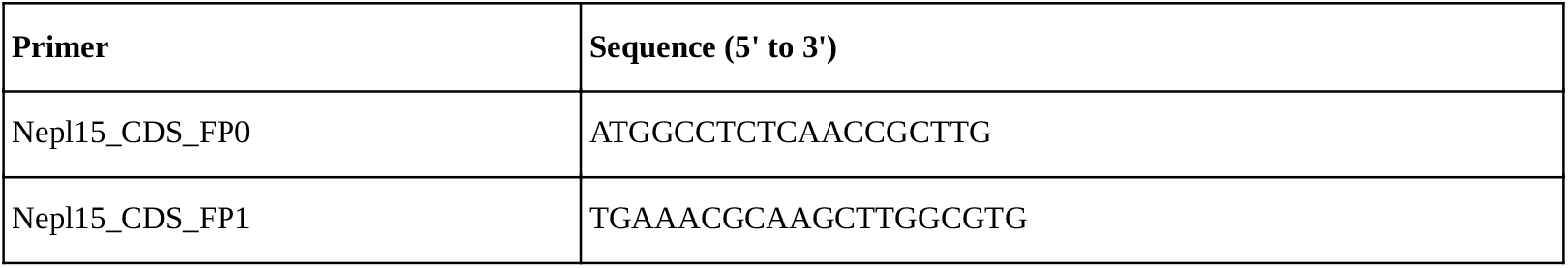
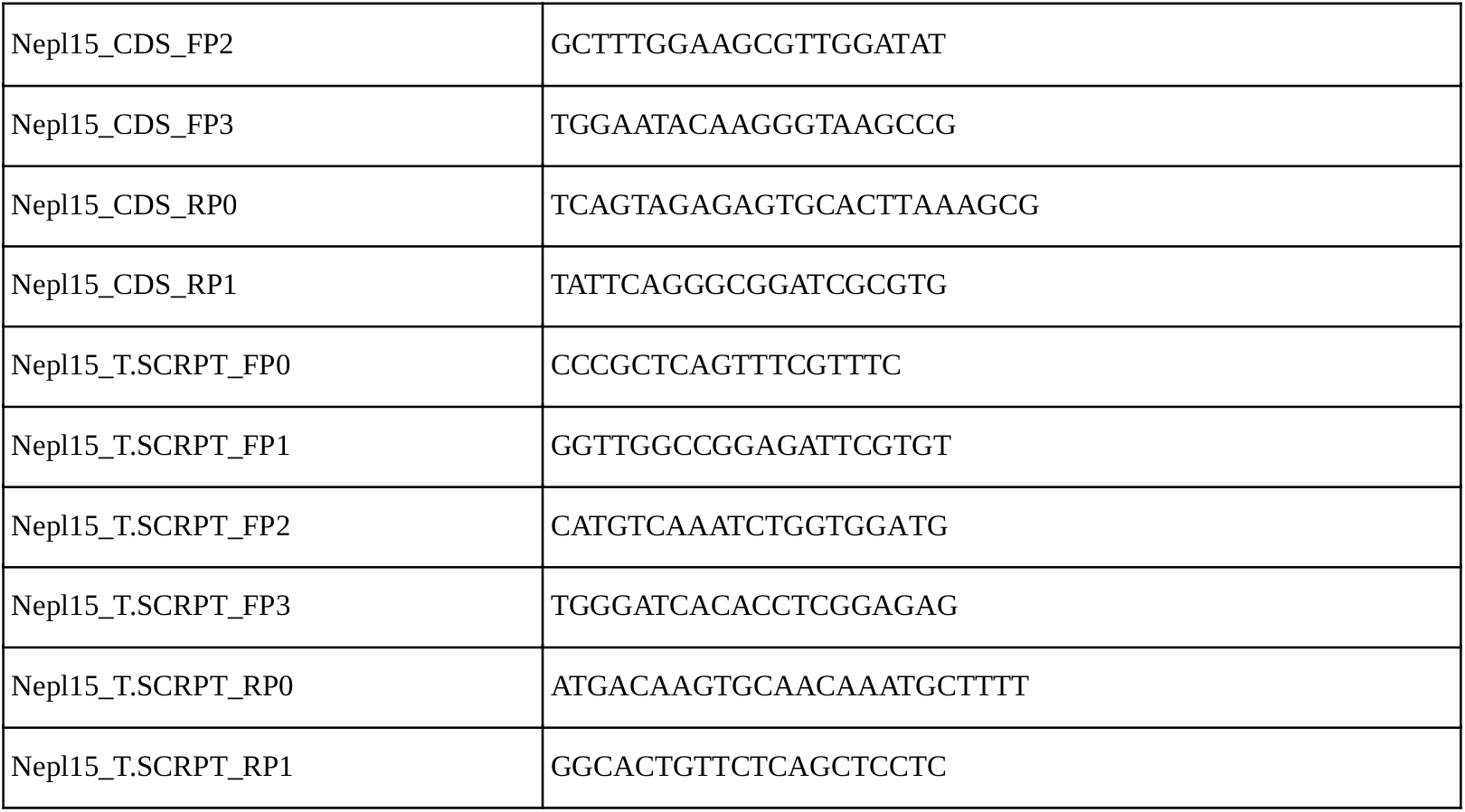
List of primers used for PCR and sequencing.

Subsequent PCR was performed using Q5® High-Fidelity 2X Master Mix (New England Biolabs-M0492S) using a custom protocol based off the protocol provided by New England Biolabs for Q5® High-Fidelity 2X Master Mix using a Bio-Rad T100™ Thermal Cycler. The PCR procedure involved the initial denaturation of cDNA at 98 °C for 3 minutes, and 30 cycles of denaturation at 98 °C for 30 seconds, annealing at 65 °C for 20 seconds, and extension at 72 °C for 1 minute and 30 seconds. The final extension occurred at 72 °C for 5 minutes and was held at 4 °C until it was ready to be loaded into a 1% agarose gel with a 1 kb ladder and stained with SYBR™ Safe DNA gel stain (Invitrogen-S33102).

### Purification and sequence alignment

The PCR products from the steps above were purified using a NucleoSpin gel and PCR clean-up kit (Macherey-Nagel-740609.50) following the manufacturer’s instructions up to the last step in which DEPC water was used to elute the DNA rather than the provided NE Buffer solution. Sanger sequencing of the PCR samples was performed by Azenta Life Sciences (https://www.genewiz.com/?lang=en) using the *Nepl15 FB CDS* or *Nepl15 FB T*.*SCRPT* primers and additional primers made about every 500 bases to avoid technical errors (Table 3). Sequence results provided various contigs that were used to create four full-length sequences which were then aligned with the original FlyBase sequence using SerialCloner (v2.6.1; http://serialbasics.free.fr/Serial_Cloner.html). Variations in codon sequences were recorded using SerialCloner’s built in BLASTn and BLASTp alignment tools and Clustal Omega (Figure 1. A) (https://www.ebi.ac.uk/jdispatcher/; retrieved 2/17/2026) (Madeira et al., 2024). Missense mutations were then noted. Translated sequences were also analyzed for changes in *Nepl15* motif regions, cleavage sites, and transmembrane helices using Clustal Omega, SerialCloner, SignalP - 6.0 (https://services.healthtech.dtu.dk/services/SignalP-6.0/; retrieved 2/13/2025) (Teufel et al., 2022), and TMHMM – 2.0 (https://services.healthtech.dtu.dk/services/TMHMM-2.0/; retrieved 2/13/2025) (Krogh et al., 2001), respectively.

## Reagents

## Supporting information

Supplemental protein prediction data

Supplemental sequencing data

## Acknowledgements

I would like to thank TTU TrUE Scholars Undergraduate Research Program, AHA SURE (Supporting Undergraduate Research Experiences) scholars’ program: Dr. Peter Keyel, Dr. Surya Banerjee’s research lab and members: Shahira Arzoo and Rubaia Tasmin.

## Author Contributions

Chase Drucker: data curation, formal analysis, funding acquisition, investigation, software, writing - original draft, visualization. Surya Banerjee: conceptualization, methodology, project administration, resources, supervision, validation, writing - review editing.

## Reviewed By

**History**

**Copyright:** © 2026 by the authors. This is an open-access article distributed under the terms of the Creative Commons Attribution 4.0 International (CC BY 4.0) License, which permits unrestricted use, distribution, and reproduction in any medium, provided the original author and source are credited.

**Citation:** Drucker C, Banerjee S. 2026. Validation of a Single *Nepl15* Transcript in Oregon-R *Drosophila melanogaster* with Minor Coding Sequence Variation Relative to FlyBase. microPublication Biology.

