## Supplementary figures and images for "Validation of a Single *Nepl15* Transcript in Oregon-R *Drosophila melanogaster* with Minor Coding Sequence Variation Relative to FlyBase"

### af686_coverage.png

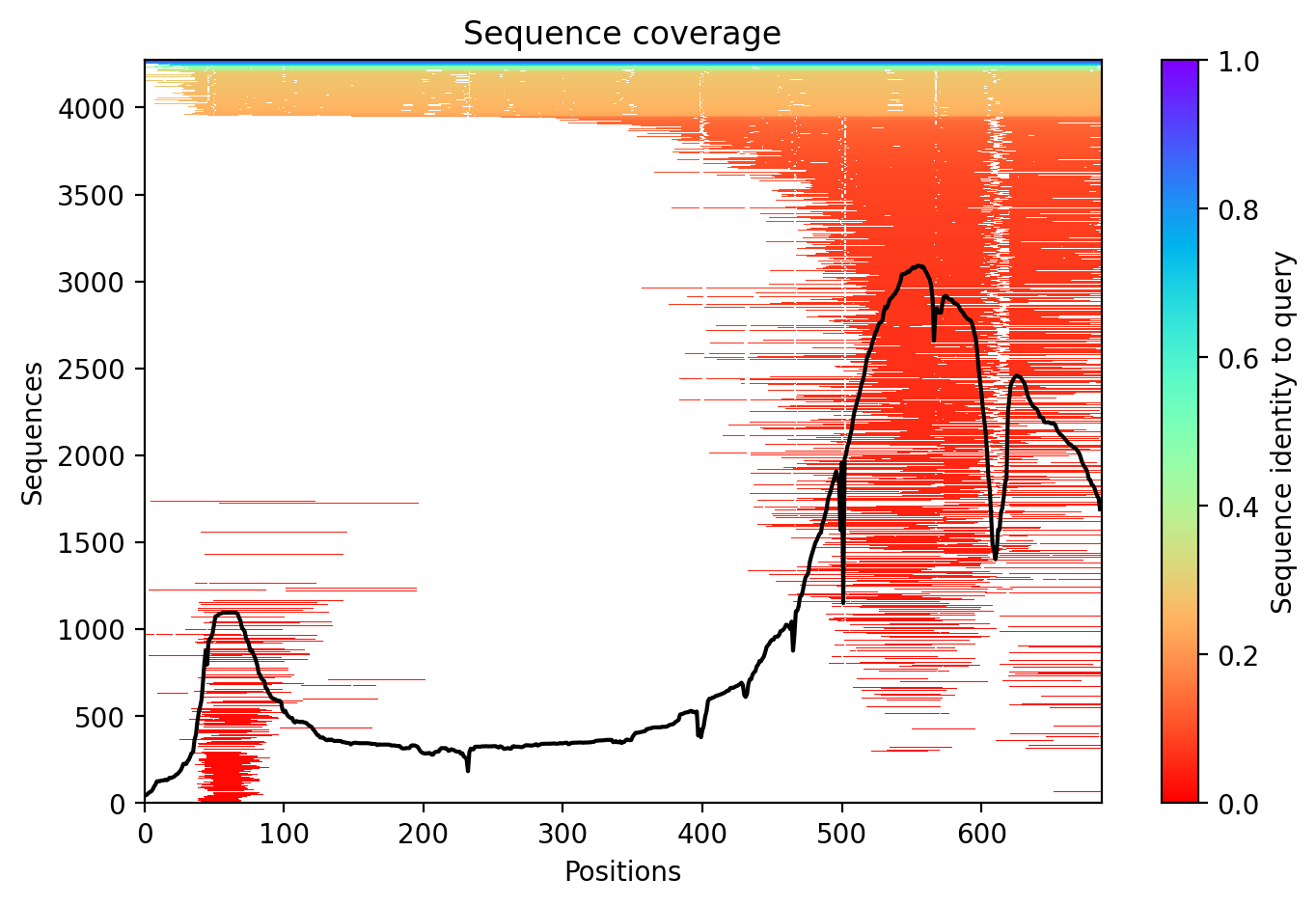

### af686_pae.png

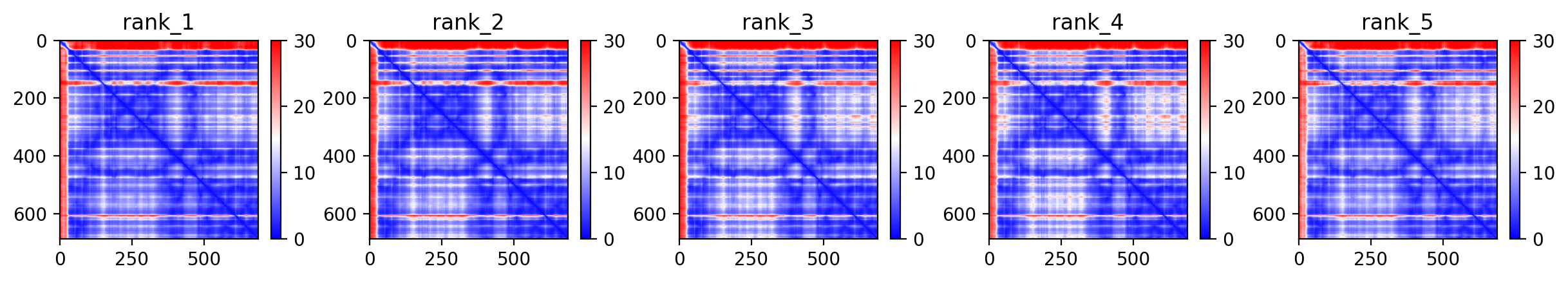

### af686_plddt.png

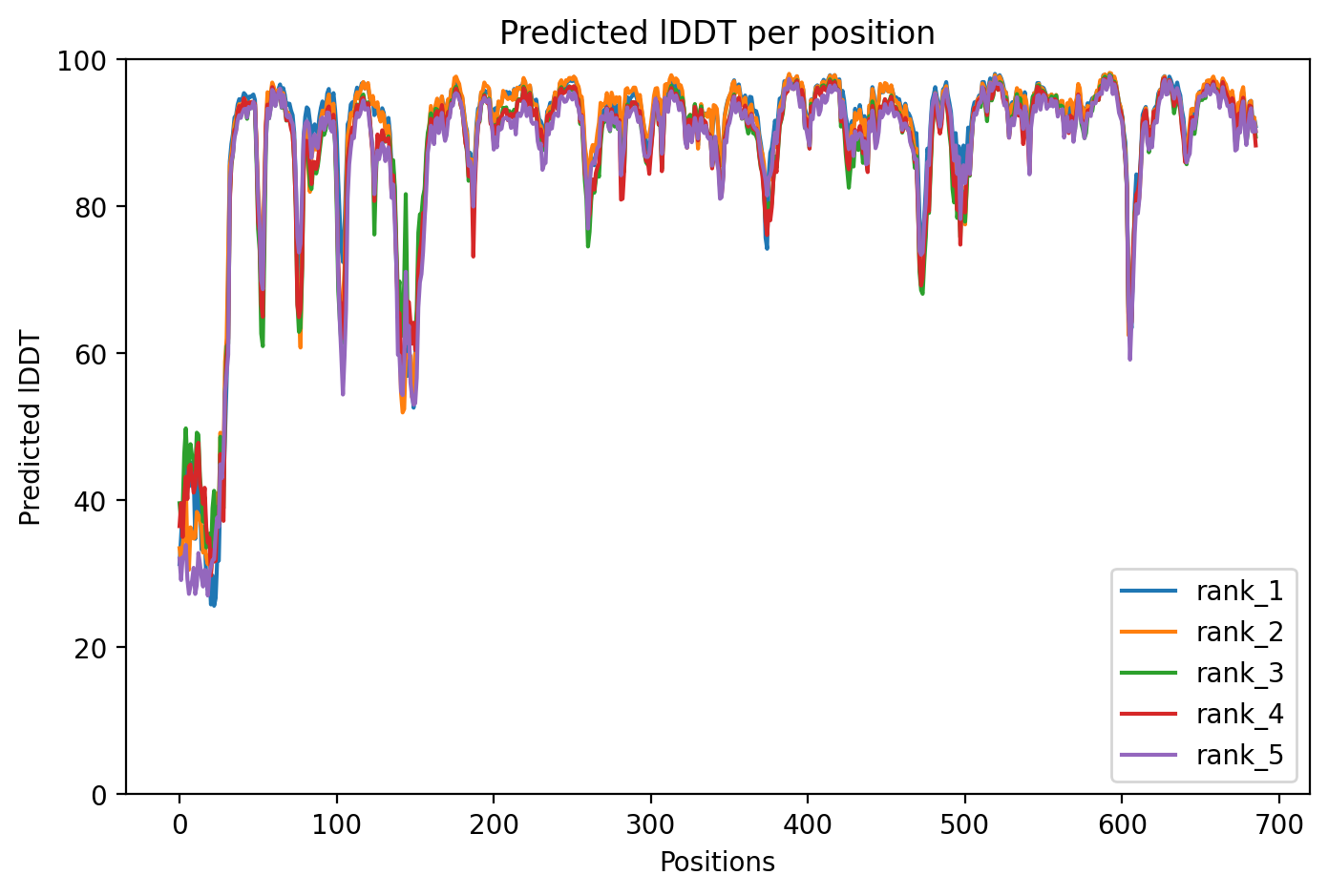
